# Viral RNA is a target for *Wolbachia*-mediated pathogen blocking

**DOI:** 10.1101/2020.04.03.023556

**Authors:** Tamanash Bhattacharya, Irene L.G. Newton, Richard W. Hardy

## Abstract

The ability of the endosymbiont *Wolbachia pipientis* to restrict RNA viruses is presently being leveraged to curb global transmission of arbovirus-induced diseases. Past studies have shown that virus replication is limited early in arthropod cells colonized by the bacterium, although it is unclear if this phenomenon is replicated in mosquito cells that first encounter viruses obtained through a vertebrate blood meal. Furthermore, these cellular events neither explain how *Wolbachia* limits dissemination of viruses between mosquito tissues, nor how it prevents transmission of infectious viruses from mosquitoes to vertebrate host. In this study, we try to address these issues using an array of mosquito cell culture models, with an additional goal being to identify a common viral target for pathogen blocking. Our results establish the viral RNA as a cellular target for *Wolbachia-* mediated inhibition, with the incoming viral RNA experiencing rapid turnover following internalization in cells. This early block in replication in mosquito cells initially infected by the virus thus consequently reduces the production of progeny viruses from these same cells. However, this is not the only contributor to pathogen blocking. We show that the presence of *Wolbachia* reduces the per-particle infectivity of progeny viruses on naïve mosquito and vertebrate cells, consequently limiting virus dissemination and transmission, respectively. Importantly, we demonstrate that this aspect of pathogen blocking is independent of any particular *Wolbachia-*host association and affects viruses belonging to *Togaviridae* and *Flaviviridae* families of RNA viruses. Finally, consistent with the idea of the viral RNA as a target, we find that the encapsidated virion RNA is less infectious for viruses produced from *Wolbachia-*colonized cells. Collectively, our findings present a common mechanism of pathogen blocking in mosquitoes that establish a link between virus inhibition in the cell to virus dissemination and transmission.

**AUTHORS SUMMARY:** Viruses transmitted by arthropod vectors pose a significant global health risk. Incidence of diseases caused by these viruses can thus be reduced by implementing effective vector control strategies. This need is further exacerbated due to the lack of commercially available vaccines and antivirals. Presence of the intracellular bacteria *Wolbachia pipientis* is associated with virus inhibition in multiple mosquito vectors. Furthermore, *Wolbachia* is inherited transovarially and spreads across the vector population like a natural gene drive, making it an attractive vector control agent. In this study, we examine how the presence of the bacterium in arthropod cells prevents initial establishment of vertebrate cell derived viruses. Our results indicate rapid turnover of incoming viral RNA very early during infection in *Wolbachia-*colonized cells, thus establishing it as a cellular target for pathogen blocking. Additionally, upon evaluating how these events might further limit virus spread, we find that infectivity of progeny viruses belonging to multiple RNA virus families are reduced on a per-particle basis. This aspect of virus inhibition is independent of any particular *Wolbachia-*host association and affects how these viruses replicate in naïve mosquito and vertebrate cells, thus providing a collective basis of reduced virus dissemination and transmission in *Wolbachia-*colonized mosquitoes.

## INTRODUCTION

The pathogen blocking ability of the arthropod endosymbiont *Wolbachia pipientis* makes it an exciting biocontrol agent that is currently being used to limit transmission of arboviruses around the world [1]. The importance of studying the underlying mechanism of pathogen blocking, however, is not only to determine the long-term feasibility of this strategy, but to also learn how this particular arthropod host-endosymbiont association enables the former to become refractory to a wide range of RNA viruses. The degree of virus inhibition varies between different *Wolbachia-*host associations and is dependent on the *Wolbachia* strain and the arthropod host species [2-5]. Collectively, three major aspects of pathogen blocking have been reported consistently across all associations: inhibition of viruses possessing positive-sense single-stranded RNA (+ ssRNA) genomes, limited virus dissemination, and lower virus transmission [2, 6-8]. Earlier studies have primarily concentrated on identifying cellular events that lead to virus inhibition. Indeed, while this approach has proven to be useful in identifying host determinants of pathogen blocking, too often results from these studies are limited to a single *Wolbachia-*host association, giving the impression that different *Wolbachia-*host permutations involve distinct mechanisms of inhibition [3, 9-12]. Furthermore, these studies do not address the question of how virus inhibition in *Wolbachia-*colonized cells ultimately limit virus dissemination in, and transmission by mosquitoes. To this end, the goal of this study was to utilize an array of mosquito cell culture models representing different *Wolbachia-*host combinations to identify a common viral target for pathogen blocking and to determine the link between intracellular virus inhibition, restricted virus dissemination in mosquito cells and virus transmission from mosquitoes to vertebrates. Our results identify the viral RNA genome as a target for pathogen blocking in arthropods, which include both fruit flies and mosquitoes. In *Wolbachia-*colonized cells, viral RNA targeting occurs at multiple stages of the replication cycle, notably in the very early stages following virus internalization and genome delivery, that lead to a shortened half-life of the incoming viral RNA. Additionally, we demonstrate that viral RNA present within viruses produced from *Wolbachia-*colonized cells are less infectious in vertebrate cells, contributing to the overall reduced infectivity of the progeny viruses. We provide evidence that this aspect of pathogen blocking is likely independent of any particular *Wolbachia* strain and arthropod host association and affects members of at least two + ssRNA virus families. Finally, we show that two major aspects of pathogen blocking i.e. limited virus dissemination and transmission occur due to the inability of these less infectious viruses to propagate in naïve arthropod and vertebrate cells.

## RESULTS

### Viral RNA is a shared target for multi-stage inhibition in *Wolbachia-*colonized cells

Presence of *Wolbachia* is associated with reduced viral gene expression in arthropod cells. This widely reported aspect of virus inhibition can be observed both *in vivo* (Figure 1A; Welch-corrected unpaired t-test, p < 0.0001, t = 3.445, df = 6.688) as well as in cell culture (Figure 1B; Welch-corrected unpaired t-test, p < 0.0001, t = 10.29, df = 6.363) [9-11, 13]. However, this quantification represents total viral RNA accumulated over multiple rounds of virus replication in mosquito cells. To determine how inhibition occurs in the initial stages of infection in the vector, we monitored the spread of vertebrate cell-derived viruses in naïve mosquito cells with and without *Wolbachia*. Spread of BHK-21-derived viruses expressing a fluorescent reporter protein (CHIKV-mKate) was thus assessed in *Aedes albopictus* mosquito cells following a synchronized infection that involved virus adsorption at 4°C. Cell monolayers were then extensively washed with 1XPBS to remove any unbound viruses and warm media (37°C) was added to cells to initialize virus internalization and infection. Virus spread was then measured over 50 hours by quantifying mean virus-encoded fluorescent reporter expression observed over four distinct fields of view taken per well every 2-hours (Supplementary Figure 1A). Two-way ANOVA was used to determine the effect of destination cell type (with or without *Wolbachia*) and/or time on virus spread. Virus growth was significantly reduced over time in cells colonized with *Wolbachia* compared to cells without the bacterium; Ordinary Two-way ANOVA, Sidak’s multiple comparisons test, *Wolbachia:* p < 0.0001, Time: p < 0.0001, Time X *Wolbachia:* p < 0.0001 (Supplementary Figure 1).

**Figure 1.**
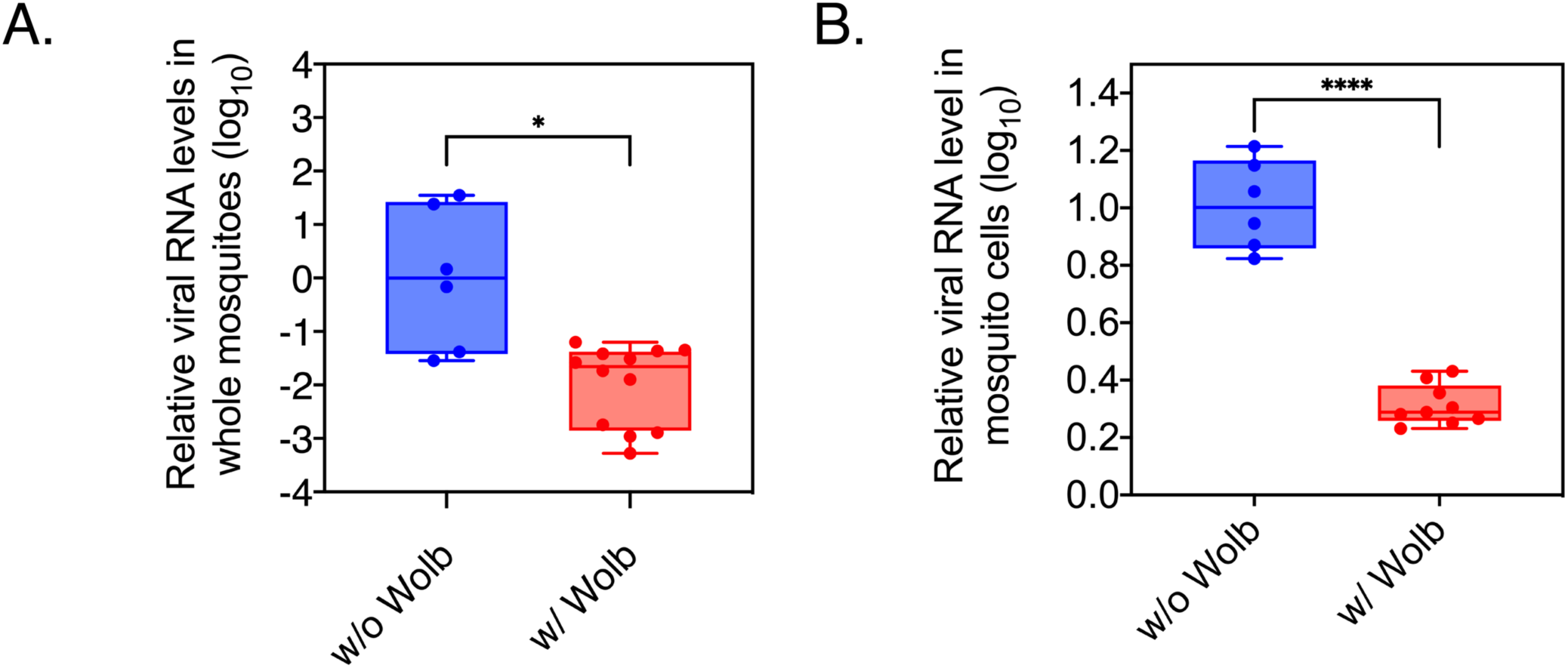
Viral RNA levels are reduced *in vivo* and in mosquito cell culture. (A) Viral RNA levels were quantified in whole female *Aedes aegypti* mosquitoes with and without *Wolbachia* (*w*AlbB strain) using qRT-PCR at 7-days post infectious blood meal with SINV. Error bars represent standard error of mean (SEM) of biological replicates (n=6). Welch’s t-test performed on log-transformed values. *P < 0.05 (B) Viral RNA replication in mosquito cells colonized with *Wolbachia*. RML12 mosquito cells with or without *Wolbachia* (*w*Mel strain) were infected with SINV at MOI of 10. Total cellular RNA was harvested forty-eight hours post infection and assayed for viral RNA levels using quantitative RT-PCR. Error bars represent standard error of mean (SEM) of biological replicates (n=6). Welch’s t-test performed on log-transformed data. ****P < 0.0001.

While our previous result validates the initial stage of virus inhibition in the vector, it is unknown how early this inhibition takes place following initial infection in the presence of *Wolbachia*. Previous studies by other groups have indicated that inhibition of both alphaviruses and flaviviruses occurs at an early stage of infection in *Wolbachia-*colonized arthropod cells [12-13]. Furthermore, data from the latter study indicate that viral RNA is susceptible to degradation immediately post-internalization [12]. Therefore, to determine whether viral RNA is degraded faster in *Wolbachia-*colonized cells following virion internalization, we asked if the incoming viral RNA half-life is altered between cells with and without *Wolbachia*. We also asked whether or not this event is exclusive to mosquito cells.

C710 *Aedes albopictus* cells with (*w*Stri) or without *Wolbachia* (w/o *w*Stri) and JW18 *Drosophila melanogaster* cells with (*w*Mel) or without *Wolbachia* (w/o *w*Mel) were grown overnight in media containing the cross-linkable nucleoside analog 4-thiouridine (4SU) (Figure 2A). In the cell, 4SU is converted to 4S-UTP before being incorporated into newly synthesized RNA, thus allowing labelling of all cellular RNA. Sindbis virus (SINV) derived from vertebrate BHK-21 cells grown in normal media, and therefore containing an unlabeled virion RNA genome, was then used to synchronously infect the 4SU-treated cells at a MOI=10. Following virus adsorption at 4°C, cell monolayers were extensively washed with 1XPBS to remove any unbound viruses. Warm media (37°C) containing 4SU was added to cells to initialize virus internalization and infection was then carried out under labelling conditions, thus allowing 4S-UTP incorporation into newly synthesized host and viral RNA, leaving the incoming viral RNA as the only unlabeled RNA species in the cell. All 4SU-labelled RNA was separated from the total RNA pool following biotinylation and streptavidin cleanup and levels of unlabeled RNA at each time point was measured relative to that present at 0 minutes post internalization. RNA half-life was extrapolated using non-linear, exponential decay model (Figure 2B). We observed steady monophasic decay of the incoming viral RNA in both mosquito and fly cells, either in the presence or absence of *Wolbachia* (Figure 2B). However, in each case mean half-life of viral RNA was reduced in *Wolbachia-*colonized cells. In mosquito cells, RNA half-life was reduced approximately 1.8-fold; One-phase decay, 115.2 minutes (w/o *w*Stri), 65.58 minutes (w/ *w*Stri). In comparison, half-life was reduced approximately 2.2-fold in fly cells; minutes (w/o *w*Mel), 24.4 minutes (w/ *w*Mel) (Figure 2C). Based on these results, we conclude that the incoming viral RNA undergoes faster turnover in *Wolbachia-* colonized cells, reducing the cellular pool of viral RNA very early in the replication cycle. Since both viral RNA synthesis and protein expression are abrogated in the presence of *Wolbachia*, our data alongside observations made by Thomas et.al, further support the idea of viral RNA as a cellular target for pathogen blocking [12].

**Figure 2:**
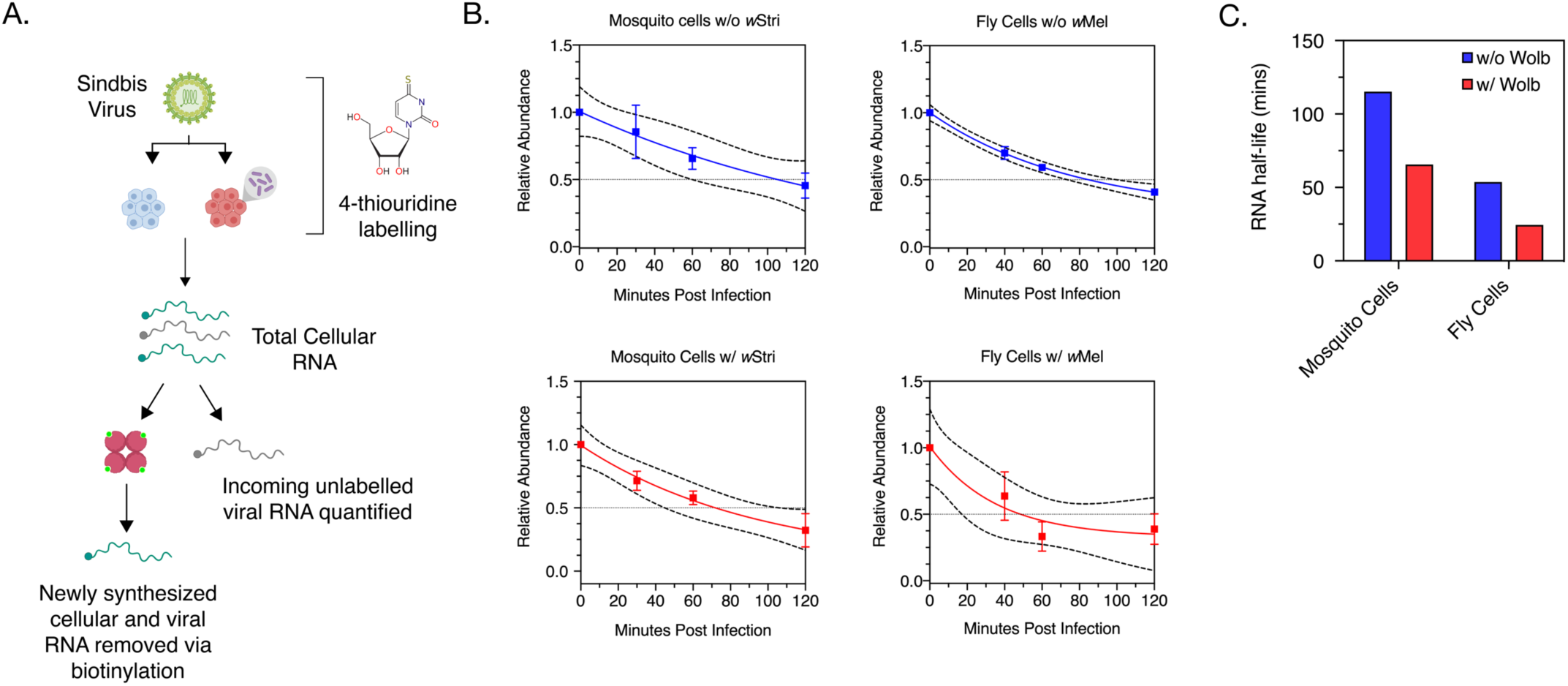
Incoming viral RNA is degraded quicker in *Wolbachia-*colonized cells. Half-life of incoming viral RNA was assessed following infection of C710 *Aedes albopictus* cells colonized with and without *Wolbachia* (*w*Stri) and JW18 *Drosophila melanogaster* cells with and without *Wolbachia* (*w*Mel). (A) Schematic representation of the experiment. Sindbis virus derived from vertebrate BHK-21 cells was used to synchronously infect C710 or JW18 cells pre-labelled with 4-thiouridine (4SU) for 12 hours. Infection was carried out under labelling conditions. At indicated times post infection, total RNA was extracted from cells, biotinylated and streptavidin beads were used to isolate incoming unlabeled viral RNA from 4SU-labelled cellular and newly synthesized RNA. Viral RNA was quantified at each time-point using quantitative RT-PCR and relative abundance was assessed relative to viral RNA detected at the start of infection (0h). (B) Relative abundance of incoming viral RNA in cells colonized with and without *Wolbachia* over 120 minutes post-infection were determined by qRT-PCR analysis as described in materials and methods. Regression analyses was performed to determine the viral RNA decay profile (represented by solid lines). Dashed lines represent the 95% CI of the aforementioned regression. (C) Viral RNA half-lives were determined from data showed in (B) using non-linear, one phase exponential decay model.

### Progeny viruses generated from *Wolbachia* colonized cells are less infectious

It is unclear how the aforementioned cellular events lead to reduced virus dissemination within the mosquito vector as well as reduced transmission into vertebrate hosts. In our previous study, we reported reduced infectivity of Sindbis viruses derived from *Wolbachia-*colonized fly cells on vertebrate BHK-21 cells [11]. In light of this result, we wondered whether these progeny viruses are compromised in their ability to infect and propagate in naïve vertebrate and arthropod cells. We reasoned that lower production of total viruses from *Wolbachia-*colonized cells in combination with their inability to propagate in naïve arthropod cells might explain why virus dissemination is reduced in mosquitoes carrying *Wolbachia*. Additionally, the observed loss in transmission could occur as a result of these viruses being unable to spread and kill vertebrate cells. But given that *Wolbachia*-mediated reduction of virus dissemination and transmission has been observed against multiple RNA viruses across multiple *Wolbachia*-host associations, we first wanted to expand our previous findings to determine whether our previous observation regarding the loss in per-particle infectivity is dependent on any particular virus or limited to only certain *Wolbachia-*host associations [2, 6-8, 11].

Viruses relevant to our study i.e. alphaviruses and flaviviruses are not known to form empty virions [14]. Thus, all virus particles produced from cells, regardless of infectivity, contain viral RNA cargo that can be measured using quantitative PCR. We therefore quantified viral genome copies present in the cell supernatant as a proxy for total virus particles released following infection in an arthropod host i.e. *Aedes albopictus-*derived or *Drosophila melanogaster-*derived cells with and without *Wolbachia*. Infectious viruses present in the same cell supernatant were assayed by quantifying plaque-forming units on vertebrate cells. Following independent quantification these attributes in tandem, we then calculated per-particle infectivity or specific infectivity ratio (SI) of progeny viruses as the ratio of infectious virus to total virus (Figure 3A).

**Figure 3.**
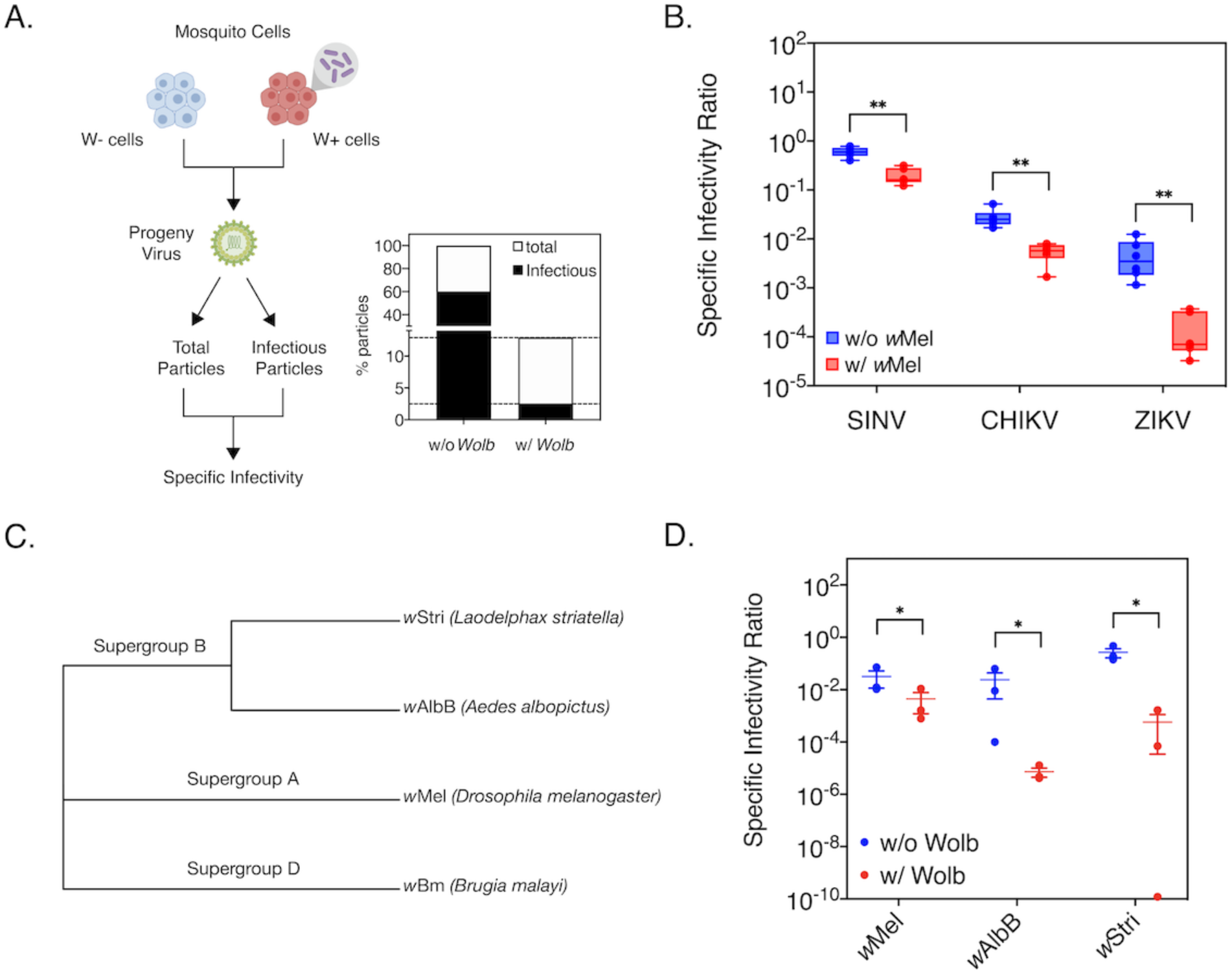
Presence of *Wolbachia* in mosquito cells reduces progeny virus infectivity in mammalian cells. RML12 mosquito cells with and without *Wolbachia* (*w*Mel) were infected with alphaviruses Sindbis (SINV), Chikungunya (CHIKV) or flavivirus Zika (ZIKV) at MOI of 10. Viral supernatants were harvested at 48 hours post infection after which infectious virus titer was quantified by performing plaque assays on BHK-21 (SINV, CHIKV) and Vero (ZIKV) cells. Total virus particles were determined by quantifying viral genome copies present in the supernatant using qRT-PCR. Reported specific infectivity (SI) ratios were calculated as total infectious virus titer divided by total particles produced per mL of viral supernatant. (A) Percentage of total SINV particles produced from cells with and without *Wolbachia* that are infectious on BHK-21 cells. (B) Specific Infectivity Ratios of progeny SINV, CHIKV and ZIKV viruses. (C) Maximum-likelihood tree representing the phylogenetic relationship between the *Wolbachia* strains used in the study was generated using MEGA X, using a MUSCLE alignment of concatenated sequences of multi-locus typing (MLST) genes (*coxA, gatB, ftsZ, hcpA, fbpA)*. Sequences from *Wolbachia* strain *w*Bm, native to the filarial nematode *Brugia malayi*, was used as a distant outgroup. Scale bar represent branch lengths. (D) Specific Infectivity of progeny SINV derived from *Aedes albopictus* cells colonized with non-native (*w*Mel and *w*Stri) and native (*w*AlbB) *Wolbachia* strains. Cells were infected with virus at MOI=0.1 and infectious virus titer produced after 48 hours was quantified via plaque assays on BHK-21 cells. Error bars represent standard error of mean (SEM) of biological replicates (n=3-6). Student’s t-test performed on log-transformed values. *P<0.05, **P < 0.01.

First, we assessed the ability of a single *Wolbachia* strain-type, *w*Mel, to inhibit multiple RNA viruses across two different arthropod host cell types, mosquito (*Aedes albopictus* RML12 cells) and the native fruit fly (*Drosophila melanogaster* JW18 cells). Mosquito cells were challenged with a panel of arboviruses, including alphaviruses Sindbis (SINV) and Chikungunya (CHIKV) and one flavivirus, Zika (ZIKV). In all cases, viruses grown in the presence of *Wolbachia* (W+ viruses) exhibited a lower SI ratio than viruses grown in cells without the endosymbiont (W-viruses). Degree to which infectivity (SI) was reduced varied depending on the virus type; SINV: 3-fold reduction (t-test, p = 0.000059, df=10, t-ratio = 6.627), CHIKV: 5-fold reduction (t-test, p = 0.000071, df=10, t-ratio = 6.6.471), ZIKV: 32-fold reduction (t-test, p = 0.000144, df=10, t-ratio = 5.934) (Figure 3B). We obtained similar results when we tested the virus panel in *Drosophila melanogaster* cells with and without *Wolbachia* (*w*Mel); SINV (t-test, p = 0.0045, df=9.458, t-ratio = 3.698), CHIKV (t-test, p = 0.013204, df=10, t-ratio = 3.006), ZIKV (t-test, p = 0.033939, df=7, t-ratio = 2.629) (Suppl. Fig 2). Presence of the same *Wolbachia* genotype in different arthropod hosts thus leads to reduced infectivity of arboviruses belonging to different virus families, indicating that this phenotype is independent of virus type and any specific *Wolbachia-*host association.

We next examined whether different *Wolbachia* strains are capable of reducing progeny virus infectivity in the context of a single arthropod host cell type. A panel of three *Aedes albopictus* derived mosquito cells were obtained, each colonized by a distinct *Wolbachia* strain, including *Wolbachia* strain *w*AlbB (Aa23 cells) derived from the native *Aedes albopictus* and non-native strains *w*Stri (C710 cells) derived from a planthopper, *Laodelphax striatellus* and the previously described *w*Mel (RML12 cells), derived from *Drosophila melanogaster* (Fig 3C) [Fallon et.al. 2012]. These cells were then challenged with SINV and progeny virus infectivity was calculated as before. We found that SI of progeny W+ SINV to be reduced in each case, irrespective of the *Wolbachia* strain present in these mosquito cells (Figure 3D; *w*AlbB (One-way ANOVA on log-transformed data with Tukey’s post-hoc test, p = 0.0257), *w*Stri (Kruskal-Wallis test on log-transformed data, p = 0.04953)). Based on these results, we therefore conclude that presence of *Wolbachia* is commonly associated with a concomitant reduction in progeny virus infectivity.

### Progeny viruses spread poorly in naïve mosquito cells

We next monitored the ability of progeny viruses derived from mosquito cells colonized with (W+ virus) and without (W-virus) *Wolbachia* to propagate in naïve mosquito cells. CHIKV expressing a fluorescent reporter protein (CHIKV-mKate) was grown in C710 *Aedes albopictus* cells with and without *Wolbachia* (*w*Stri), purified and subsequently used to infect naïve mosquito cells at equal MOIs (MOI=5) following a synchronous infection. Virus infection and spread was then monitored over a period of 48 hours using a live-cell imaging system by quantifying mean virus-encoded fluorescent reporter expression observed over four distinct fields of view taken per well every 2-hours (Figure 4A). Three-way ANOVA was used to determine the effects of virus source (Source i.e. W- or W+ virus), destination cell type (with or without *Wolbachia*) and/or time, on virus spread. Our results show significant effects of all three variables on virus growth, both on their own as well as in combination with each other; Three-way ANOVA, Tukey’s multiple comparisons test, Source: p < 0.0001, *Wolbachia*: p < 0.0001, Time: p < 0.0001, Source X Time: p < 0.0001, Source X *Wolbachia*: p < 0.0001, *Wolbachia* X Time: p < 0.0001, Source X Time X *Wolbachia*: p = 0.0016 (Figure 4B). As expected, W-virus grew more poorly in naïve cells with *Wolbachia* compared to those without (Figure 4B, square symbols). However, W+ viruses (Figure 4B, circle symbols) exhibited reduced growth in naïve cells both in the presence and absence of *Wolbachia*. Importantly, these results indicate that reduced infectivity of W+ viruses might contribute to their inability to spread to new mosquito cells regardless of whether or not these cells are colonized by *Wolbachia*. Assuming that the mechanism of cellular inhibition of virus replication is cell-autonomous, this model might help explain why limited virus dissemination occurs within tissues with both high and low *Wolbachia* densities. Our results are supported by the observation made previously by Dutra et.al, where viruses isolated from salivary gland secretions of *Wolbachia-*colonized mosquitoes failed to establish systemic infections in naïve mosquitoes following injection [6].

**Figure 4.**
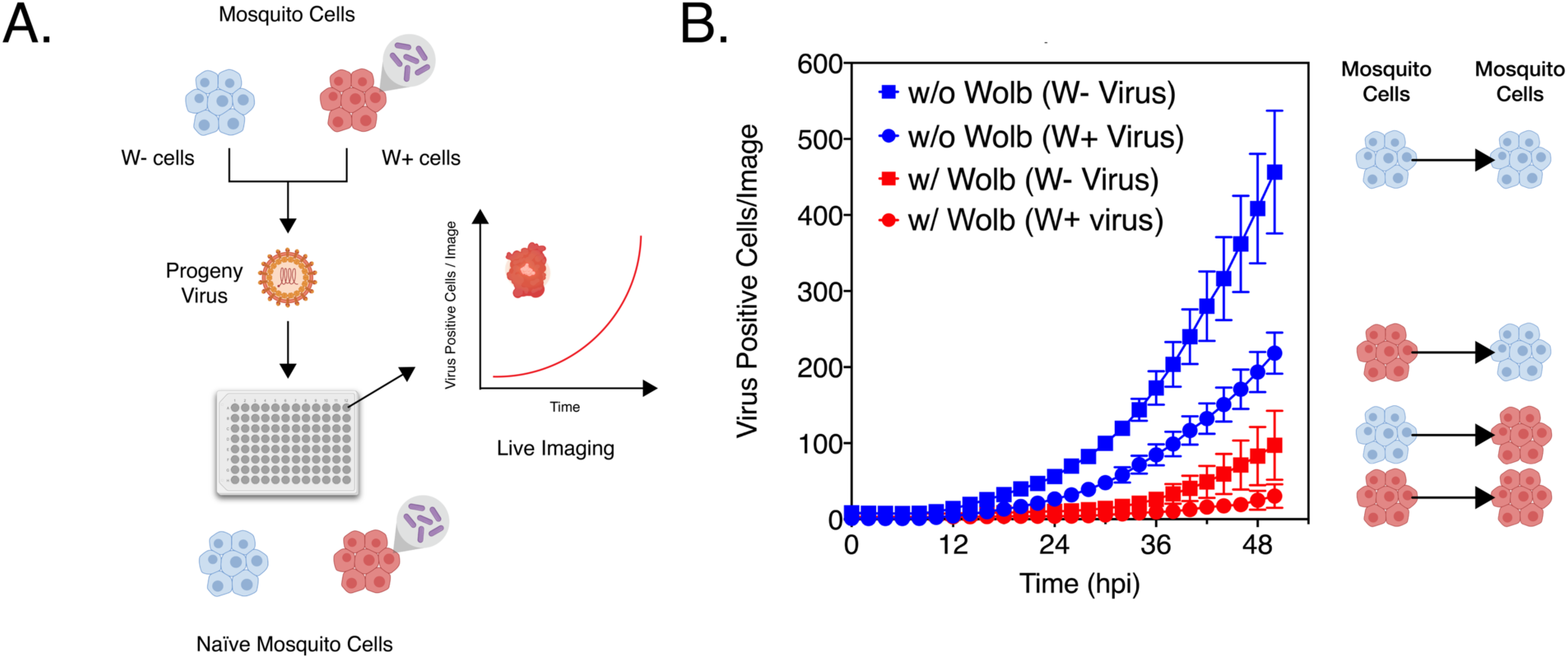
Progeny viruses derived from *Wolbachia* colonized cells replicate poorly in naïve mosquito. (A) Schematic representation of the experiment. CHIKV expressing mKate fluorescent protein from a second sub-genomic promoter was grown in C710 *Aedes albopictus* cells in the presence (W+ virus) or absence (W- virus) of *Wolbachia* (*w*Stri strain). These progeny viruses were then used to infect naïve C710 cells with (depicted in red) and without (depicted in blue) *Wolbachia* synchronously at a MOI=1 particle/cell. Virus growth in cells, plated on a ninety-six-well plate, was measured in real time by imaging and quantifying the number of red cells (Virus Positive Cells/Image) expressing the virus encoded mKate protein over a period of forty-eight hours, using live cell imaging. (B) Color of the data points distinguish the two destination cell lines where virus replication was assayed on; blue represent C710 cells without *Wolbachia* while red represent C710 cells with *Wolbachia*. Shape of the data points refer to the nature of the progeny viruses used to initiate infection; squares represent W- viruses, grown in C710 cells without *Wolbachia*, while circles represent W+ viruses, grown in C710 cells with *Wolbachia*. Y-axis label (Virus Positive Cells/Image) represent red cells expressing virus-encoded mKate fluorescent protein in a single field of view, four of which were averaged/sample at every two-hour time point collected over the course of infection. Three-way ANOVA of multivariate comparisons. Error bars represent standard error of mean (SEM) of biological replicates (n=6).

### Progeny viruses spread poorly in naïve vertebrate cells

We further assessed the infectivity of W+ progeny viruses to spread in vertebrate cells using live-cell imaging to validate our earlier results (Figure 3B). As before, purified progeny W+ and W-viruses derived from mosquito cells colonized with (W+ virus) and without (W-virus) *Wolbachia* (*w*Stri) was subsequently used to infect naïve BHK-21 cells at a MOI of 5 particles/cell. Infection was synchronized as before and virus spread was measured over 42 hours by quantifying mean virus-encoded fluorescent reporter expression observed over four distinct fields of view taken per well every 2-hours (Figure 5A). Two-way ANOVA was used to determine the effect of virus source (Source i.e. W- or W+ virus) and/or time on virus spread. Spread of W-viruses were consistently faster compared to W+ viruses, with peak number of W-virus positive cells observed at 26 hours post infection compared to 30 hours post infection for W+ viruses; Ordinary Two-way ANOVA, Source: p = 0.0261, Time: p < 0.0001, Time X Source: p = 0.0068 (Figure 5B). Interestingly, peak number of virus positive cells was higher for W+ viruses relative to W-viruses between 28 and 42 hours. However, this is due to a delay in cell death in W+ infected cell populations, thus allowing greater and prolonged expression of virus encoded reporter. In comparison, cells infected with W-viruses succumb early to infection, resulting in a faster loss in reporter activity between 26 and 42 hours (Figure 5B).

**Figure 5:**
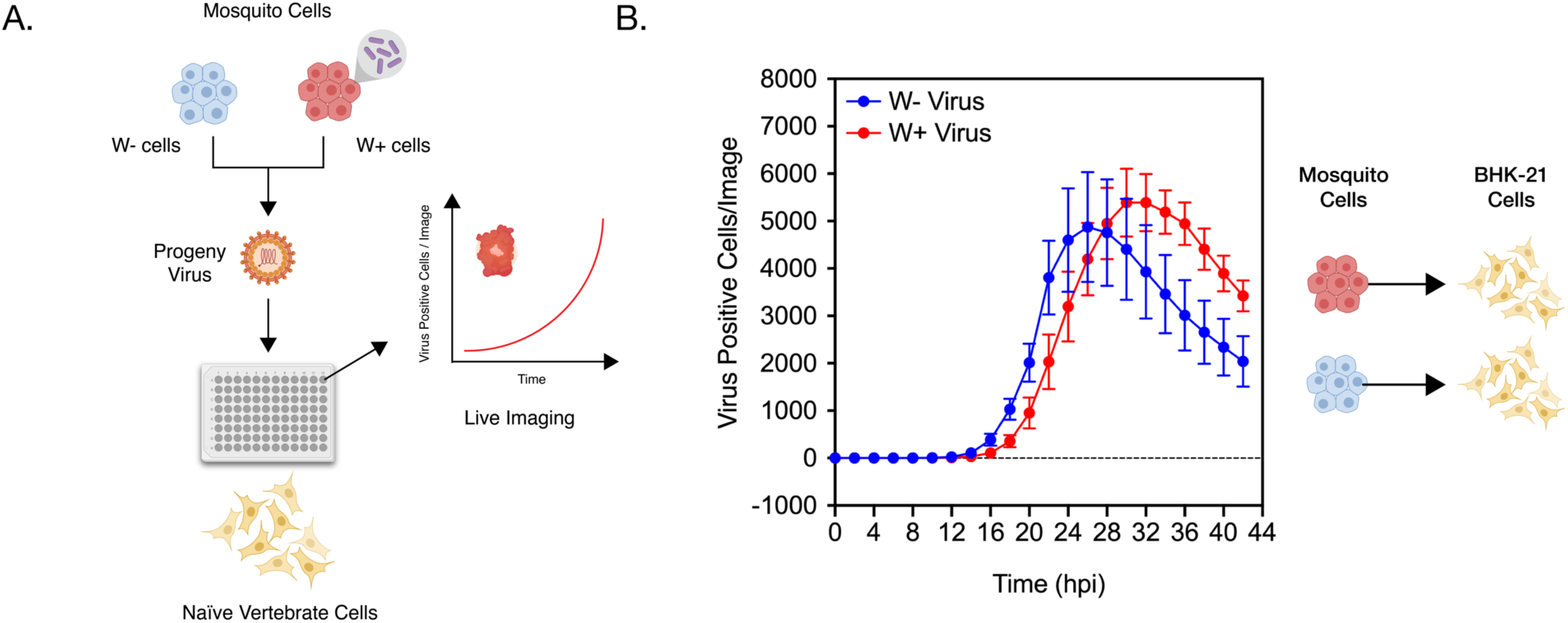
Progeny viruses derived from *Wolbachia* colonized cells replicate poorly in naïve vertebrate cells. (A) Schematic representation of the experiment. CHIKV expressing mKate fluorescent protein from a second sub-genomic promoter was grown in C710 *Aedes albopictus* cells in the presence (W+ virus) or absence (W- virus) of *Wolbachia* (*w*Stri strain). These progeny viruses were then used to infect naïve vertebrate BHK-21 cells synchronously at a MOI=5 particles/cell. Virus growth in cells, plated on a ninety-six-well plate, was measured in real time by imaging and quantifying the number of red cells expressing the virus encoded mKate protein over a period of forty-two hours using live-cell imaging. (B) Color of the data points distinguish the progeny viruses used to initiate infection in BHK-cells; blue represent progeny viruses derived from C710 cells without *Wolbachia* (W- virus) while red represent progeny viruses derived from C710 cells with *Wolbachia* (W+ virus). Y-axis label (Virus Positive Cells/Image) represent red cells expressing virus-encoded mKate fluorescent protein in a single field of view, four of which were averaged/sample at every two-hour time point collected over the course of infection. Two-way ANOVA of multivariate comparisons. Error bars represent standard error of mean (SEM) of biological replicates (n=6).

### Encapsidated viral RNA within progeny viruses are less infectious

While reduced infectivity of progeny viruses explains limited virus dissemination and transmission in *Wolbachia*-colonized mosquitoes, it is not evident from our previous data why W+ viruses are less infectious. Simplistically, two key factors can result in the observed loss in virus infectivity; compromised virion structure and dysfunctional viral RNA. The former might impair virus attachment and entry to cells, while the latter would prevent efficient virus replication post entry. We reasoned that if W+ viruses exhibit structural defects that impair their ability to bind and enter cells, direct delivery of the encapsidated viral RNA into the cell should bypass this blockade and thus allow genome replication comparable to W-virus derived RNA. Therefore, we isolated encapsidated virion RNA from W+ and W-viruses produced from *w*Mel-colonized mosquito cells and assessed their ability to replicate in naïve vertebrate BHK-21 cells following transfection. For this replication assay, we used SINV with nanoluciferase reporter fused to the nonstructural open reading frame (SINV-nLuc) and measured luciferase activity as a proxy for viral non-structural protein expression over time (Fig 6A). First, we used progeny viruses to a establish a synchronized infection in BHK-21 cells as described earlier and measured viral replication over a period of 9 hours post infection. We found W+ virus replication to be significantly reduced relative to W-viruses over time, reaffirming our earlier results regarding poor infectivity of W+ viruses on vertebrate cells; Two-way ANOVA of multivariate comparisons, Time: p < 0.0001, *Wolbachia*: p < 0.0001, Time X *Wolbachia*, p < 0.0001 (Fig 6B). Next, we isolated virion encapsidated RNA from W+ and W-viruses and transfected naïve BHK-21 cells with equal viral genome copies (Fig 6A). As before, we used luciferase reporter activity as a proxy for virus replication over time and found significantly reduced reporter activity in cells transfected with W+ virus-derived RNA (Fig 6C). This reduction in reporter activity was more severe compared to the reduction observed in our previous experiments where infections were initiated with W+ viruses (Fig 6B). We have previously demonstrated that interactions between Sindbis virus capsid protein and the viral RNA (SINV C: R interaction) are important in regulating the function of the incoming viral RNA in vertebrate cells [15]. Absence of critical SINV C: R interaction sites that impact association of the viral RNA with the capsid proteins lead to increased degradation of the incoming viral RNA, reducing the half-life by almost 2.5 - fold. As SINV C: R interactions are absent during our transfection experiments, it is possible that virion RNA derived from W+ viruses are turned over at a faster rate, causing the observed reduction in replication of W+ virus-derived RNA (Fig 6C). Finally, to test whether reduced replication of W+ virion RNA result in the production of fewer infectious units, we quantified plaque-forming units following transfection of virion RNA into BHK-21 cells and found W+ virion RNA to produce 10-times fewer infectious units relative to W-virion RNA after 48 hours (Fig 6D). Taken together, these results suggest that encapsidated viral RNA present within W+ viruses are deficient in their ability to replicate in naïve vertebrate cells, ultimately resulting in the formation of fewer infectious units.

**Figure 6.**
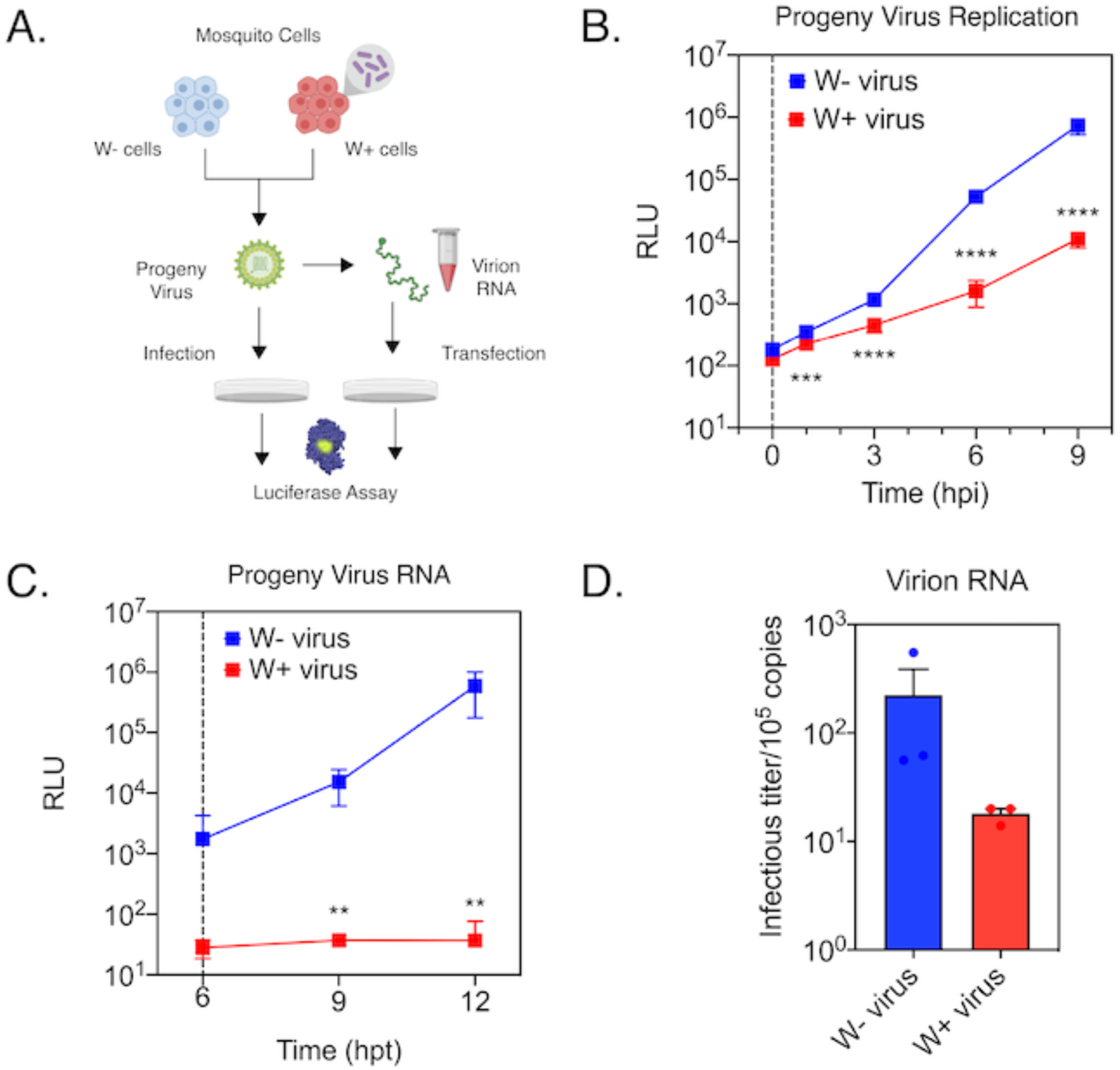
RNA encapsidated within progeny viruses derived from *Wolbachia* colonized cells are less ctious. (A) Sindbis viruses expressing nanoluciferase reporter gene (SINV-nLuc) were derived from RML12 mosquito cells with (W+ virus) or without (W- virus) *Wolbachia* (*w*Mel) and were subsequently used to synchronously infect naïve BHK-21 cells at equivalent MOIs of 5 particles/cell. Cell lysates were collected at indicated times post infection and luciferase activity (RLU) was measured and used as a proxy to quantify viral replication. (B) Approximately 10^5^ copies each of virion encapsidated RNA extracted from the aforementioned W+ and W- viruses were transfected into naïve BHK-21 cells and infectious titer was determined by the counting the number of plaques produced after 48 hours post transfection. (C) Replication kinetics of the previously isolated virion encapsidated RNA was determined by measuring luciferase activity following transfection into BHK-21 cells (RLU) as before. Error bars represent standard error of mean (SEM) of biological replicates (n=3). Student’s t-test performed on log-transformed values. **P<0.01, ***P < 0.001, ****P < 0.0001.

Considering that viral RNA and protein synthesis is abrogated in the presence of *Wolbachia*, we conclude based on the data presented above that the viral RNA serves as a target for endosymbiont-mediated inhibition in arthropod cells.

## DISCUSSION

Uptake of viruses occur as the mosquito takes an infectious bloodmeal from a vertebrate animal. As the blood meal is digested, these viruses infect midgut cells before escaping the midgut barrier and disseminating to other mosquito tissues, which become persistently infected. Therefore, it is important to note that the viruses initially establishing infection in the vector are of vertebrate origin, while those that undergo dissemination and eventually transmission, are derived from mosquito cells. Prior studies have shown that initial infection of viruses in mosquito midguts is reduced in the presence of *Wolbachia*, suggesting an inability of vertebrate-derived viruses to establish infection [7, 17]. In this study, we show that presence of *Wolbachia* restricts vertebrate cell-derived virus growth in naïve mosquito cells. While past studies have shown inhibition of viral RNA and protein synthesis in *Wolbachia-*colonized cells, it is still unclear how early virus inhibition occurs in cells that are initially infected [12-13]. Using Semilki Forest Viruses (SFV) carrying a translationally fused luciferase reporter, Rainey et.al. demonstrated reduced expression of reporter genes at 7 hours post infection in fly cells colonized with *Wolbachia*, suggesting that inhibition occurs early in the infection process [13]. Importantly, the authors also observed reduced luciferase expression following transfection of an *in vitro* transcribed virus replicon reporter, which indicates that inhibition of virus replication is independent of virus entry into *Wolbachia-*colonized cells and therefore might involve an intracellular virus target [12-13]. More recently, Thomas et.al. reported that *Wolbachia-* mediated inhibition of mosquito cell (C6/36) derived DENV-2 in *w*Mel-colonized Aag-2 cells involve degradation of the viral RNA [12]. Interestingly, the authors observe reduced abundance of viral RNA in Aag2-*w*Mel cells as early as 1h post infection, following synchronized binding (4°C) and internalization (25°C) of DENV-2 viruses. It is important to note that this reduction likely represent a loss of incoming viral RNA, due to low amounts of nascently synthesized viral RNA being present in the cell this early in infection. Here using metabolic labelling, we determined the half-life of the incoming vertebrate cell-derived viral RNA to show that virus inhibition occurs very early during infection, with the incoming viral RNA undergoing faster turnover in *Wolbachia-*colonized cells. This likely involves one or more cellular RNA degradation pathways, including perhaps orthologs of the RNA exonuclease *Xrn1*, whose role in pathogen blocking of flaviviruses has been demonstrated in *Aedes aegypti* derived Aag2-*w*Mel cells, further suggesting a common mechanism of action across different cell types and RNA viruses [12].

Presence of *Wolbachia* in mosquitoes has been shown to also reduce the rate of virus infection in different mosquito tissues. These include tissues proximal to the midgut like Malpighian tubules and fat bodies, as well as distal tissues like salivary glands, implying limited virus dissemination in the presence of the endosymbiont [7]. Interestingly, the degree to which virus inhibition occurs in these tissues is not correlated with either *Wolbachia-*density or its role in innate immunity. In fact, viral RNA levels are dramatically reduced in the mosquito head, which is poorly colonized by *Wolbachia* [7]. These results are confounding, given that the mechanism of pathogen blocking is thought to be cell-autonomous [18]. In addition, virus transmission is demonstrably reduced in *Wolbachia-* colonized mosquitoes [2, 6-8]. It is important to note that in this case, viruses transmitted to a vertebrate animal following a secondary bloodmeal are produced in the mosquito salivary gland tissues. In *Wolbachia-*colonized mosquitoes, reduced transmission is thought to likely occur as a result of reduced virus growth in salivary gland tissues. Interestingly, however, previous studies have found that although salivary gland secretions from *Wolbachia-*colonized mosquitoes contain detectable levels of viral genome copies, these levels far exceed the actual number of infectious viruses that can be assayed on either vertebrate or mosquito cells, implying a reduction in per-particle infectivity [6]. Similar results have also been obtained in cell culture [19].

We have previously shown reduced per-particle infectivity of Sindbis virus grown in *Drosophila melanogaster* cells colonized with *w*Mel on vertebrate BHK-21 cells [11]. In this study, we provide evidence that this attribute of pathogen blocking is present in *Aedes albopictus* mosquito cells, colonized with both native (*w*AlbB) and non-native (*w*Mel and *w*Stri) *Wolbachia* strains. Additionally, *Wolbachia* reduces the infectivity of progeny viruses belonging to two distinct families of + ssRNA viruses, *Togaviridae* (SINV, CHIKV) and *Flaviviridae* (ZIKV) in both mosquito and fly cells. Using growth assays in vertebrate cells, we clearly demonstrate the inability of progeny W+ viruses to replicate in naïve vertebrate cells, thus linking virus infectivity to loss in transmission. Importantly, we demonstrate that reduced infectivity of these progeny viruses also limits their ability to grow in naïve mosquito cells. Remarkably, limited virus growth is independent of whether or not these naïve cells are colonized with *Wolbachia*, thus providing an explanation for why viral RNA levels are reduced in tissues with low *Wolbachia* titers. While further investigation is required to determine whether these results are limited to single virus type and/or host cell species, we expect this phenotype to remain consistent in *Wolbachia-* host associations where pathogen blocking have been reported thus far.

Finally, we demonstrate that loss in virus infectivity occur at the level of the encapsidated virion RNA and affects its ability to replicate following direct delivery into vertebrate cells. Further work is required to determine the exact fate of W+ virus derived viral RNA in vertebrate cells in terms of its stability, localization and translation. Given the absence of information regarding potential structural differences between viruses derived from mosquito cells with and without *Wolbachia*, our data does not exclude the possibility of an additional block in progeny virus binding that might exacerbate this inhibitory effect.

Notably, our findings support the hypothesis of viral RNA being the target for pathogen blocking. *Wolbachia’s* ability to restrict viruses is seemingly limited to positive-sense single-stranded RNA (+ssRNA) viruses, as inhibition of neither negative-sense single-stranded RNA (-ssRNA) nor DNA viruses have been observed either in the field, or under laboratory conditions. It is therefore likely that factor(s) regulating virus inhibition affect this particular genomic feature shared by all susceptible viruses [3, 20-21]. We have previously reported the role of *Drosophila melanogaster* RNA methyltransferase Dnmt2 as a host determinant of pathogen blocking [11]. Interestingly, the mosquito ortholog of Dnmt2 has also been shown to play an important role in pathogen blocking in mosquitoes, albeit in a manner opposite to flies [22]. Furthermore, loss of Dnmt2 levels in fly cells significantly increase viral RNA replication and progeny virus infectivity, implying an antiviral mechanism of action that might involve targeting of the viral RNA. Given the biological role of Dnmt2 as an RNA cytosine methyltransferase, future studies will also focus on exploring the possibility of changes in the methylation profile of viral RNA in the presence and absence of *Wolbachia*.

## MATERIALS AND METHODS

### Insect and Mammalian Cell Culture

RML12 *Aedes albopictus* cells with and without *Wolbachia w*Mel (a generous gift from Dr. Seth Bordenstein, Vanderbilt University) were grown at 24 °C in Schneider’s insect media (Sigma-Aldrich) supplemented with 10% heat-inactivated fetal bovine serum (Corning), 1% each of L-Glutamine (Corning), non-essential amino acids (Corning) and penicillin-streptomycin-antimycotic (Corning). C710 *Aedes albopictus* cells with and without *Wolbachia* (a generous gift from Dr. Horacio Frydman, Boston University) were grown in 1X Minimal Essential Medium (Corning) supplemented with 5% heat-inactivated fetal bovine serum (Corning), 1% each of L-Glutamine (Corning), non-essential amino acids (Corning) and penicillin-streptomycin-antimycotic (Corning). Aa23 *Aedes albopictus* cells with and without *Wolbachia w*AlbB (a generous gift from Dr. Horacio Frydman, Boston University) were grown at 24 °C in Schneider’s insect media (Sigma-Aldrich) supplemented with 20% heat-inactivated fetal bovine serum (Corning), 1% each of L-Glutamine (Corning), non-essential amino acids (Corning) and penicillin-streptomycin-antimycotic (Corning). JW18 *Drosophila melanogaster* cells with and without *Wolbachia w*Mel were grown at 24 °C in Shields and Sang M3 insect media (Sigma-Aldrich) supplemented with 10% heat-inactivated fetal bovine serum, 1% each of L-Glutamine (Corning), non-essential amino acids (Corning) and penicillin-streptomycin-antimycotic (Corning). *Aedes albopictus* C636 cells and mammalian BHK-21 and Vero cells were grown at either 28°C (C6/36) or 37 °C (BHK/Vero) under 5% CO_2_ in 1X Minimal Essential Medium (Corning) supplemented with 10% heat-inactivated fetal bovine serum (Corning), 1% each of L-Glutamine (Corning), non-essential amino acids (Corning) and penicillin-streptomycin-antimycotic (Corning).

### Virus infection in cells and progeny virus production

Virus stocks were generated from *Aedes albopictus* derived cells (RML12, C710) or *Drosophila melanogaster* derived cells (JW18) with or without *Wolbachia* by infecting naïve cells with virus at a MOI of 10. In all cases, serum-free media was used for downstream virus purification. Media containing virus was collected 5 days post-infection for alphaviruses SINV (SINV-nLuc), CHIKV (CHIKV18125-capsid-mKate), and 7 days post-infection for flavivirus ZIKV (MR76 Uganda Strain). Virus stocks were subsequently purified and concentrated by ultracentrifugation (43K for 2.5 h) over a 27% (w/v) sucrose cushion dissolved in HNE buffer. Viral pellets were stored and aliquoted in HNE buffer before being used for all subsequent experiments. Viral titers were determined using standard plaque assay on vertebrate BHK-21 cells and virus particles were determined by quantifying viral genome copies via quantitative RT-PCR using primers listed in the primer table (Supplementary Table 1).

### Mosquito rearing and infectious blood meals

*Aedes aegypti* mosquitoes either -infected and -uninfected with *Wolbachia* strain *w*AlbB (generously provided by Dr. Zhiyong Xi, Michigan State University, USA), were reared in an insect incubator (Percival Model I-36VL, Perry, IA, USA) at 28 °C and 75% humidity with 12 h light/dark cycle. Four to six-day old mated female mosquitoes were allowed to feed for 1h on approximately 10^8^ PFUs of SINV (TE12-untagged) containing citrated rabbit blood (Fisher Scientific DRB030) supplemented with 1mM ATP (VWR) and 10% sucrose using a Hemotek artificial blood feeding system (Hemotek, UK) maintained under constant temperature of 37 °C. Engorged mosquitoes were then isolated and reared at 28 °C in the presence of male mosquitoes. Infected mosquitoes were either harvested whole or dissected at 5-7 days post blood meal using the following method. At specified time points, mosquitoes were anesthetized following a short 5 min exposure to cold, before they were transferred to a CO_2_ pad for dissections. Dissected salivary glands were collected in sterile 1XPBS (Phosphate Buffered Saline) before being snap frozen in liquid nitrogen and storage at −80 °C for further processing. Samples for qRT-PCR were homogenized in TRiZOL (Sigma Aldrich) reagent and further processed for RNA extractions.

### Virion RNA extraction and transfection

Virion encapsidated RNA was extracted from viruses (SINV-nLuc) were purified over a 27% sucrose cushion using TRiZOL reagent (Sigma Aldrich) using manufacturer’s protocol. Post extraction, RNAs were DNase (RQ1 RNase-free DNase, NEB) treated using manufacturer’s protocol to remove cellular contaminants and viral RNA copies were quantified using quantitative RT-PCR using primers probing for SINV nsP1 and E1 genomic regions (Primer Table). To determine infectivity or replication kinetics of virion derived RNA, equal copies of viral RNA or equal mass of virion derived total RNA, quantified using qRT-PCR, were transfected into BHK-21 cells for SINV in serum-free Opti-MEM (Gibco). Transfection was carried out for 6 hours before the transfection inoculum was removed and overlay was applied. Cells were fixed 48 hours post transfection for SINV using 10% (v/v) formaldehyde and stained with crystal violet to visualize plaques.

### Live cell imaging

Growth of fluorescent reporter viruses in cells were monitored using IncuCyte live cell analyses system (Essen Biosciences, USA). Cells were grown under standard conditions as described earlier under 5% ambient CO_2_ either at 37°C for vertebrate BHK-21 cells and 27°C for *Aedes albopictus* C710 cells. Cells were plated to 75-80% confluency in 96-well plates to allow distinct separation between adjacent cells and preserve cell shape for optimal automated cell counting. Cells per well were imaged and averaged across four distinct fields of view, each placed in one quarter of the well, every two hours over the course of the infection. For every sample, total fluorescence generated by cells expressing the red fluorescent reporter mKate was calculated and normalized by the cell number. A manual threshold was set to minimize background signal via automated background correction at the time of data collection.

### Luciferase Based Viral Replication Assays

Quantification of viral genome and sub-genome translation was performed using cellular lysates following synchronized infections with reporter viruses (SINV-nLuc), or transfections with virion-derived RNA from the aforementioned viruses. At indicated times post infection, samples were collected and homogenized in 1X Cell Culture Lysis Reagent (Promega). Samples were mixed with NanoGlo luciferase reagent (Promega), incubated at room temperature for 3 minutes before luminescence was recorded using a Synergy H1 microplate reader (BioTech instruments).

### Real-time quantitative RT-PCR analyses

Following total RNA extraction using TRiZOL reagent, cDNA was synthesized using MMuLV Reverse Transcriptase (NEB) with random hexamer primers (Integrated DNA Technologies). Negative (no RT) controls were performed for each target. Quantitative RT-PCR analyses were performed using Brilliant III SYBR green QPCR master mix (Bioline) with gene-specific primers according to the manufacturer’s protocol and with the Applied Bioscience StepOnePlus qRT-PCR machine (Life Technologies). The expression levels were normalized to the endogenous 18S rRNA expression using the delta-delta comparative threshold method (ΔΔCT). Fold changes were determined using the comparative threshold cycle (CT) method (Primer Table).

### Statistical analyses of experimental data

All statistical tests were conducted using GraphPad Prism 8. The average fold change (FC) in qRT-PCR experiments was calculated using the variable bootstrapping method, measuring the fold change between each potential pair of experimental and the control samples to determine the variability of the mean [23].

## ACKNOWLEDGEMENTS

We thank all members of the Hardy, Newton, Danthi, Mukhopadhyay and Patton Labs for fostering ideas through helpful discussions and for sharing their instruments. We thank our colleagues outside of IU for their generosity in sharing reagents, including Dr. Horacio Frydman, Boston University for providing us Aa23 and C710 *Aedes albopictus* cells, Dr. Seth Bordenstein, Vanderbilt University for providing us RML12 *Aedes albopictus* cells and finally, Dr. Zhiyong Xi, Michigan State University for providing us *Aedes aegypti* mosquitoes.

---

**Supplementary Figure 1.**
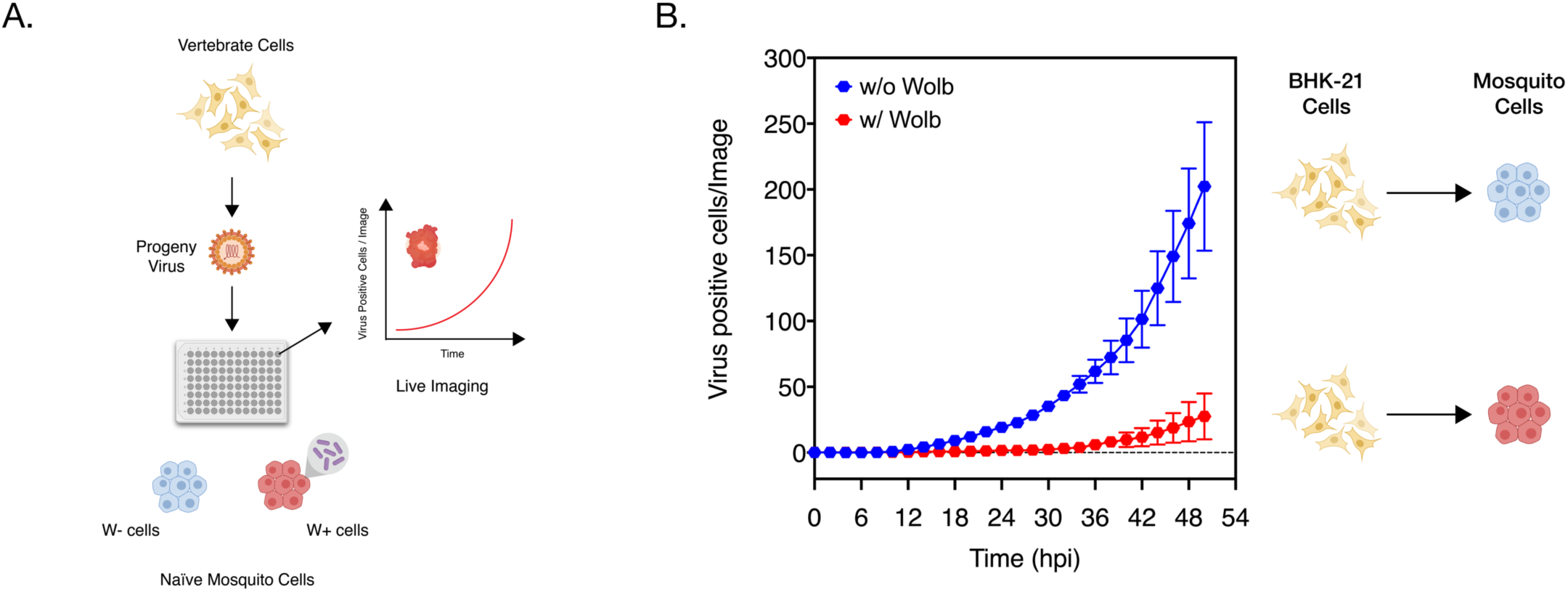
Presence of *Wolbachia* reduces spread of vertebrate-derived viruses in naïve mosquito cells. (A) Schematic representation of the experiment. CHIKV expressing mKate fluorescent protein from a second sub-genomic promoter was grown in BHK-21 cells. These progeny viruses were then used to infect naïve C710 cells with (depicted in red) and without (depicted in blue) *Wolbachia* synchronously at a MOI=1 particle/cell. Virus growth in cells, plated on a ninety-six-well plate, was measured in real time by imaging and quantifying the number of red cells (Virus Positive Cells/Image) expressing the virus encoded mKate protein over a period of forty-eight hours, using live cell imaging. (B) Color of the data points distinguish the two destination cell lines where virus replication was assayed on; blue represent C710 cells without *Wolbachia* while red represent C710 cells with *Wolbachia*. Y-axis label (Virus Positive Cells/Image) represent red cells expressing virus-encoded mKate fluorescent protein in a single field of view, four of which were averaged/sample at every two-hour time point collected over the course of infection. Two-way ANOVA of multivariate comparisons. Error bars represent standard error of mean (SEM) of biological replicates (n=7-9).

**Supplementary Figure 2.**
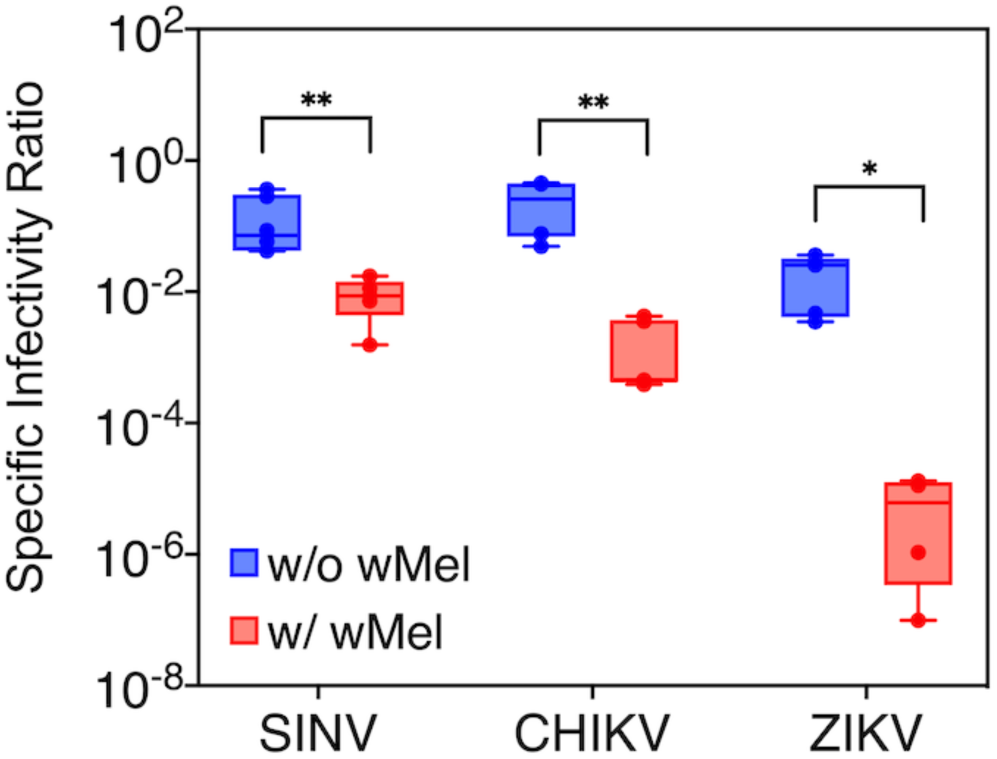
Specific Infectivity Ratios of progeny RNA viruses Sindbis (SINV), Chikungunya (CHIKV) and Zika (ZIKV) derived from *Drosophila melanogaster* JW18 cells colonized with the native *Wolbachia* strain *w*Mel. Error bars represent standard error of mean (SEM) of biological replicates (n=5-6). *P < 0.05; **P < 0.01.

**Supplementary Table 1:**
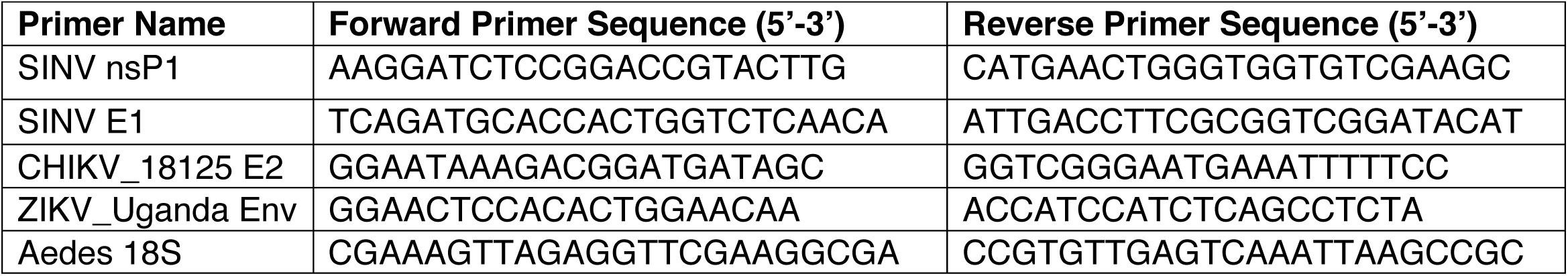
Primers used in this study. Primers were purchased from Integrated DNA Technologies (IDT). All primers were used at a final concentration of 10μM for quantitative RT-PCR reactions.

